# Melanoma cells release DEL-1 via small extracellular vesicles

**DOI:** 10.1101/2022.04.04.487024

**Authors:** Marianne Schnödl, Felix Tuchmann, Judith Wenzina, Karin Neumüller, Goran Mitulović, Marion Gröger, Gerwin Heller, Peter Petzelbauer

## Abstract

DEL-1 (developmental endothelial locus-1) induces integrin signaling and recognizes phosphatidylserine exposed on apoptotic cells. We show that DEL-1, which is thought to be a secreted molecule, is not found in melanoma cell culture supernatants but is exported by an endosomal pathway and released via small extracellular vesicles (sEV). Proteomics of DEL-1 positive sEV, but not of DEL-1 negative sEV contain proteins associated with poor survival in cancer. To determine whether DEL-1 is suitable to predict treatment responses, we isolated sEV from plasma of melanoma patients before and 90 days of treatment with checkpoint inhibitors. Although we could not detect DEL-1 in plasma sEV even in patients with progressive disease (most likely due to the very low protein yield from sEV isolated from 1ml plasma), the principal component analysis allowed a clear differentiation between controls and patients as well as between patients before and after treatment. Interestingly, in one patient with complete regression, in the post treatment sample, the protein expression profile remained in the pre-treatment cluster. The low protein yield of patient sEV and the low patient number are clear limitations of this study, but results demonstrate that this method could have the potential to predict treatment responses.

## Introduction

DEL-1 (developmental endothelial locus-1), also called EDIL3, is a glycoprotein secreted by many cell types. It induces integrin signaling by binding to the RGD sequence of integrins (1) and recognizes phosphatidylserine exposed on apoptotic cells (2). The function of DEL-1 depends on its expression site and interferes with myelopoiesis, it reduces leukocyte recruitment, promotes efferocytosis and the resolution of inflammation (3). In cancer DEL-1 promotes epithelial mesenchymal transition (4) and is an independent prognostic factor for overall survival in breast and lung cancer patients (5, 6). DEL-1 detected in small extracellular vesicles (sEV) isolated from plasma has been identified as a biomarker for early breast cancer detection (7). In melanoma cells DEL-1 is a marker for invasive behavior and high mRNA expression in melanoma samples is associated with poor patient survival (8). However, the anti-inflammatory function of DEL-1 complicates the interpretation of effects of DEL-1 in cancer as shown in a mouse model. DEL-1 deficiency in mice implanted with B16F10 melanoma cells favors melanoma lung metastasis due to reduced neutrophil accumulation, IL-17A up-regulation and reduced NK cell infiltration (9). However, this study analyzed the effect of host-derived DEL-1 and not that of melanoma-derived DEL-1. We asked DEL-1 mRNA expression in melanoma cells correlated with protein production and if and how DEL-1 is released. Moreover, we asked if this could be used to predict treatment responses in melanoma patients.

## Materials and Methods

### Cell culture

The patient-derived melanoma cell lines VM-8, VM-14, VM-15, VM-21, VM-28, VM-47, VM-48 and MCM1DLN have been described previously (8, 10-13). VM-15, VM-21 and MCM1DLN cells were grown in Dulbecco’s modified Eagle medium (DMEM) and VM-8, VM-14, VM-28, VM-47 and VM-48 cells in RPMI 1640 medium supplemented with 10% heat-inactivated fetal calf serum (FCS), 2mM L-glutamine and 50 U/ml streptomycin-penicillin (all from Gibco, Thermo Fisher Scientific, Waltham, MA, USA). HEK293T cells were provided by Martin Bilban, Medical University of Vienna. Human umbilical vein endothelial cells (HUVEC) were isolated and used as described previously (14), according to the ethics committee permission EK 1621/2020 of the Medical University of Vienna. Cells were maintained at 37°C in a humidified atmosphere containing 5% CO_2_ and regularly tested for Mycoplasma contamination.

### Isolation of extracellular vesicles

Melanoma cells were grown to 70% confluency, washed and then cultured in FCS-free medium for 4 days. Cell culture medium was harvested and centrifuged at 400xg for 5 minutes to pellet non-adherent cells, followed by centrifugation of the supernatant for 10 minutes at 1,500xg to remove remaining cellular debris. The obtained supernatant was collected and transferred to ultracentrifuge tubes (PC Oak Ridge centrifuge tubes 25×94mm) (Thermo Fisher Scientific, Waltham, MA, USA). Samples were centrifuged using a Sorvall WX ultracentrifuge and T-1250 rotor (both from Thermo Fisher Scientific, Waltham, MA, USA) for 120 minutes at 100,000xg to pellet extracellular vesicles. To separate the different fractions of extracellular vesicles, a differential ultracentrifugation protocol was used. The supernatant was first centrifuged for 30 minutes at 18,000xg to pellet microvesicles. To isolate exosomes, the remaining supernatant was transferred to a fresh ultracentrifuge tube and centrifuged for 120 minutes at 100,000xg. Pellets were resuspended in PBS and stored at -80°C for subsequent analysis. Protein content-based EV characterization according to the consensus paper (15) revealed that after the 2-step centrifugation protocol melanoma cell-derived sEV contain most category 1 proteins required for EV characterization (CD63, CD81, CD82, CD47, MHC class I, LAMP1/2, SDC*, EMMPRIN, ADAM10, NT5E, CD55, CD59, TSPAN8, CD37, CD53, CD9, CD90 and most category 2 proteins (TSG101, CHMP*, PDCD6IP, VPSA/B, ARRDC1, FLO1/2, EHD*, RHOA, ANXA*. HSP90AB1, ARF6, SDCBP). Co-isolated non-EV-related proteins are APOA1/2. Proteins associated with other intracellular compartments are LMNA, CANX, HSPA5, ATG9A, ACTN1/4, KRT18 (The complete proteome is published in ProteomeXchange with the identifier PXD025475. For the reviewer: Username; PW: UVmO0YGs).

Isolation of extracellular vesicles from human plasma was approved by the medical ethical committee of the Medical University of Vienna (ECS 1780/2018) and in accordance with the International Code of Medical Ethics of the World Medical Association (Declaration of Helsinki). Blood was collected in EDTA-coated tubes from 5 healthy volunteers (3 male and 2 female, mean age 34+15) and from melanoma patients pre (day 0) and post (day 90) immunotherapy treatment after informed consent. Detailed patient data can be found in Table 1. Plasma was isolated by centrifugation at 1,500xg for 10 minutes, aliquoted and stored at - 80°C or in liquid nitrogen. To isolate exosomes, 1ml of plasma was diluted in PBS and centrifuged at 1,500xg for 15 minutes to deplete platelets. The obtained supernatant was collected, transferred to ultracentrifuge tubes and centrifuged for 30 minutes at 18,000xg to remove microvesicles, followed by 120 minutes at 100,000xg to pellet exosomes, as described earlier. Pellets were dissolved in PBS and stored at -80°C. Protein content-based EV characterization according to the consensus paper (15) revealed that after the 2-step centrifugation protocol plasma-derived EVs contain some category 1 proteins required for EV characterization (ADAM10, CD59, CD9, GYPA) and no two category 2 proteins. Co-isolated non-EV-related proteins are APOA1/2 and ALB. No proteins associated with other intracellular compartments were detectable (The complete proteome is published in ProteomeXchange with the identifier PXD025475. For the reviewer: Username; PW: UVmO0YGs). Of note, the low protein yield of the plasma samples is certainly a limitation of this study, however, given lack of contaminating proteins supports the relative purity of the isolation.

**Table 1.**

| Melanoma patients |  |  |  |  |  | Day -1 |  |  |  |  | Day 90 |  |  |  |  |  | New metastasis at day 90 |
| --- | --- | --- | --- | --- | --- | --- | --- | --- | --- | --- | --- | --- | --- | --- | --- | --- | --- |
| Pat. | Gen. | Age | Primum | Mutation | Therapy | AJCC Stage | S100 | LDH U/L | PMN G/L | Ly G/L | AJCC Stage | Response | S100 | LDH | PMN G/L | Ly G/L |  |
| P1 | m | 55 | pectoral | NRAS | Nivo | IIIC | 0.02 | 168 | 3.4 | 1.8 | IIIC | stable | 0.02 | 179 | 3.2 | 1.9 | no new lesions |
| P2 | m | 75 | gluteal | NRAS | Pembro | IV | 0.09 | 216 | 4.9 | 2.0 | IV | progress | 0.09 | 272 | 6.5 | 2.0 | adrenal gland |
| P3 | m | 71 | plantar | N/A | Nivo | IIIC | 0.03 | 143 | 8.9 | 2.24 | IV | progress | 0.05 | 153 | 6.6 | 1.7 | inguinal |
| P4 | m | 72 | scapular | BRAF V600E | Nivo | IIIC | 0.04 | 194 | 9.3 | 2.0 | IV | progress | 0.06 | 186 | 7.9 | 2.1 | pulmonal & cutaneous |
| P5 | m | 69 | dorsal | BRAF V600E | Nivo & Pembro | IV | 0.32 | 813 | 8.0 | 1.7 | IV | response | 0.05 | 140 | 3.4 | 2.7 | complete regression |
| P6 | w | 56 | capillitial | N/A | Nivo | IIIC | 0.07 | 209 | 2.2 | 1.8 | IV | progress | 0.09 | 189 | 2.8 | 1.6 | cervical & pulmonal |
| P7 | m | 51 | oesophag eal | cKIT | Nivo & Pembro | IV | 0.05 | 178 | 4.5 | 1.1 | IV | progress | 0.06 | 167 | 2.7 | 1.0 | pulmonal & hepatal |

Pat, patient ID; Gen, gender; AJCC, American Joint Committee on Cancer; LDH, lactate dehydrogenase; PMN, polymorphonuclear leukocyte; Ly, lymphocytes; N/A, not available.

### Antibodies and reagents

Primary antibodies used in this study include the following: anti-EDIL3 (SC-293337) (Santa Cruz, Dallas, TX, USA) and anti-EEA1 (#2411) (Cell Signaling Technology, Danvers, MA, USA). The secondary antibodies used for Western Blot analysis were IRDye800CW (925-32213) and IRDye680RD (925-68072) (both from LI-COR Biosciences, Lincoln, NE, USA). Secondary antibody used for immunofluorescence was goat anti-rabbit IgG-Alexa546 (#A-11071) from Thermo Fisher Scientific (Waltham, MA, USA).

### Immunoblotting

For Western Blot analysis, whole cell and total exosome lysates were isolated using RIPA buffer (50mM Tris, 150mM NaCl, 1% NP-40, 0.5% sodium deoxycholate, 0.1% SDS), supplemented with 1% protease inhibitor and phosphatase inhibitor cocktail 2 and 3 (all from Sigma Aldrich, St. Louis, MO, USA). Samples were centrifuged at 14,000 rpm for 20 minutes at 4°C. Protein content of the supernatant was measured using Bradford Protein Assay (Bio-Rad Laboratories, Hercules, CA, USA). For whole cell lysates, a total of 30µg protein was incubated with reducing 6x Laemmli sample buffer and analyzed on a 10% SDS-PAGE gel. Gels were run at 100V for 1.5 hours. Proteins were then transferred onto a nitrocellulose membrane (GE-Healthcare, Chicago, IL, USA) for 1 hour at 15V using the Trans-Blot semi-dry transfer system (Bio-Rad Laboratories, Hercules, CA, USA). After transfer, membranes were incubated with Ponceau S staining solution (SERVA Electrophoresis, Heidelberg, Germany) to check for successful protein transfer, followed by blocking for 1 hour at room temperature with 4% skim milk powder (Sigma-Aldrich, St. Louis, MO, USA) in Tris-buffered saline pH 7.4 (TBS) with 0.1% Tween20 (TBS-T) (Bio-Rad Laboratories, Hercules, CA, USA). After one washing step with TBS-T, membranes were incubated with primary antibodies diluted in 4% bovine serum albumin (BSA) (Sigma-Aldrich, St. Louis, MO, USA) in TBS-T overnight at 4°C. The following day and upon several washing steps with TBS-T, membranes were incubated with secondary antibodies diluted in 4% milk in TBS-T for 1 hour at room temperature. After washing, blots were visualized using Odyssey CLx and analyzed with Image Studio Software Version 5.2 (both from LI-COR Biosciences, Lincoln, NE, USA).

### Transduction and transfection

For stable overexpression of DEL-1, a mammalian lentiviral gene expression system was employed. The plasmid for DEL-1 overexpression (VB900000-0562grx) was purchased from VectorBuilder (Chicago, IL, USA) and the packaging plasmid psPAX2 (#12260) and envelope plasmid pMD2.G (#12259) were obtained from Addgene (Watertown, MA, USA). The plasmid GFP-pLVTHM (#12247) (Addgene, Watertown, MA, USA) was used as a negative control. HEK293T were grown at 75% confluency in 6cm culture dishes in RPMI medium without antibiotics. One hour before transfection, HEK293T cells were treated with 25µM chloroquine. Co-transfection of the plasmids was then performed using Lipofectamine 2000 (Invitrogen, Thermo Fisher Scientific, Waltham, MA, USA) according to the manufacturers protocol with 10µg DEL-1 or control gene expression plasmid, 1.75µg pMD2.G envelope plasmid and 5.25µg psPAX2 packaging plasmid. After 6 hours, medium was changed to fresh antibiotics-free RPMI medium without chloroquine. Conditioned medium containing lentiviral vectors was collected 24, 48 and 72 hours post transfection, filtered and supplemented with 6µg/ml polybrene (Sigma-Aldrich, St. Louis, MO, USA) before it was added to MCM1DLN cells, which were seeded in T-25 flasks at 60-70% confluency. After 72 hours cells were sorted for GFP expression and analyzed for DEL-1 mRNA and protein expression.

For cellular protein localization, a construct coding for green fluorescent protein tagged DEL-1 (VB200505-128ave) was ordered from VectorBuilder (Chicago, IL, USA) and transiently expressed in VM-14 cells. Melanoma cells were seeded at 80% confluency, washed with RPMI medium without supplements and transfected in Opti-MEM (Gibco, Themo Fisher Scientific, Waltham, MA, USA) by using Lipofectamine 2000 (Invitrogen, Thermo Fisher Scientific, Waltham, MA, USA) according to the suppliers recommendations. 48 to 72 hours later, DEL-1-GFP expression was analyzed by qPCR, Western Blotting and immunofluorescence.

### Immunofluorescence

For immunofluorescence, a total of 1,500 VM-14 cells per well were seeded in ibidi P-Slide 8 well chambered coverslips (ibidi, Gräfelfing, Germany) and transiently transfected with a construct coding for green fluorescent protein tagged DEL-1 (VB200505-128ave) (VectorBuilder, Chicaco, IL, USA) as described previously. Cells were fixed with 4% paraformaldehyde in PBS for 10 minutes at room temperature, washed three times with PBS and permeabilized with permeabilization solution (20mM Hepes, 300mM sucrose, 50mM NaCl, 0.5% Triton X-100) for 5 minutes at -20°C. After washing with PBS, the primary antibody was diluted in PBS with 1% BSA and incubated over night at 4°C. After additional washing steps, the secondary antibody diluted in PBS with 1% BSA was added for 1.5 hours at room temperature. Slides were imaged with LSM 780 confocal microscope with AiryScan detector. Images were processed using ArivisVision 4D software (Arivis, Munich, Germany) to segment the exosomes. This was performed with the Blob Finder followed by area measurement and event counting.

### Electric cell-substrate impedance sensing (ECIS) measurement

After equilibration of 8W10E+ ECIS array plates (ibidi, Gräfelfing, Germany) with medium for 16 hours at 37°C, HUVEC were seeded on the gelatin-coated electrode chambers at a density of 12,000 cells/well in 300µl of medium. Impedance at multiple frequencies ranging from 62.5 to 64,000 Hz was then continuously monitored using the ECIS Z instrument (Applied BioPhysics, Troy, NY, USA) as described earlier (16). Upon reaching a stable plateau at 4,000Hz, PBS, recombinant human DEL-1 protein (6046-ED) (R&D Systems, Minneapolis, MN, USA) and/or RGD peptide (A8052) (Sigma-Aldrich, St. Louis, MO, USA) were added to the cells. Transendothelial resistance change at 250 Hz was analyzed by using the ECIS software Version 1.2.186 (Applied BioPhysics, Troy, NY, USA).

### Mass spectrometry analysis

Exosomes were isolated as described previously and dissolved in PBS. For exosomes derived from melanoma cell lines, low molecular-weight contaminant removal was performed using Exosome spin columns MW3000 (Thermo Fisher Scientific, Waltham, MA, USA), according to the manufacturers protocol. This step was omitted for human plasma exosome samples. Subsequently, all exosome samples were diluted with one volume of Exosome Resuspension Buffer and one volume of 2X Denaturing Solution (both from Invitrogen, Thermo Fisher Scientific, Waltham, MA, USA), followed by protein precipitation with dichloromethane (Merck, Darmstadt, Germany) and methanol (Thermo Fisher Scientific, Waltham, MA, USA). Briefly, three volumes of methanol, one volume of dichloromethane and three volumes of distilled water were added to one sample volume and incubated at -20°C for 20 minutes. Then, samples were centrifuged at 9,000xg for 5 minutes at room temperature. The upper phase was discarded and three volumes of methanol were added. Samples were centrifuged at 9,000xg for 10 minutes and the supernatant was discarded again after which pellets were air-dried. Dry pellets were dissolved in 50µl 50mM triethylammoniumbicarbonate (Sigma Aldrich, St. Louis, MO, USA) containing 0.1% RapiGest SF (Waters, Milford, MA, USA). Protein concentration was measured by a nano spectrophotometer (DS-11, DeNovix, Wilmington, NC, USA). An aliquot of 20µg protein was further reduced using 5mM dithiothreitol for 30 minutes at 60°C and alkylated for 30 minutes in the dark by using 15mM iodoacetamide (both from Sigma-Aldrich, St. Louis, MO, USA). Digestion with MS grade porcine trypsin (1:20, w/w) (Thermo Fisher Scientific, Waltham, MA, USA) was carried out overnight at 37°C and aliquots of 1µg digested protein were prepared and stored at -80°C until injection.

The nano chromatographic separation was performed using a nano-RSLC UltiMate 3000 system (Thermo Fischer Scientific, Waltham, MA, USA). For sample loading and desalting, a PepMap C18trap-column (Thermo Fischer Scientific, Waltham, MA, USA) of 300 µm ID x 5 mm length, 5 µm particle size, 100 Å pore size was used. Nano HPLC Separation of digested proteins was performed using a 200 cm C18 µPAC separation column (PharmaFluidics, Ghent, Belgium). The dimensions of the µPAC column were: 5 µm pillar diameter, 2.5 µm inter-pillar distance, 18 µm pillar height; 315 µm bed channel width. The pillars are superficially porous, end-capped with C18-chains.

The aqueous loading mobile phase contained 2% ACN,0.1% TFA, and 0.01% HFBA cooled to 3°C. The mobile phase transferred the sample from the sample loop to the trap column by the loading pump at 30 µl/min to the trapping column, which was operated in the column oven at 50°C. The trapping time was set to 10 min, which was sufficient to load the sample to the trapping column and wash out possible salts and contaminants. The cooled mobile phase enabled trapping of the hydrophilic analytes at enhanced oven temperature. Peptides were separated and analyzed using positive nano HPLC-ESI-MS. The gradient elution was performed by mixing two mobile phases composed of the following solvents:

- Mobile phase A (MPA): 95% H2O, 5% ACN, 0.1% FA;
- Mobile phase B (MPB): 50% ACN, 30% MeOH, 10% TFE (2,2,2-Trifluoro ethanol), 10% - 0.1% FA.

At 200 minutes, the trapping column was switched back into the separation flow and equilibrated for the following run.

Due to the long separation path, the void volume of these columns is higher than conventionally packed nano separation columns. Therefore, the flow rate of the nano HPLC pump was set to 800 nl/min for the first 10 minutes with gradual lowering to 600 nl/min for 11 minutes for separation purposes. The gradual increase of the flow rate at the end of the separation run to 800 nl/min is performed for speeding the equilibration of the separation column. Details on developing and testing different gradients for the 200-cm µPAC column are described by (17).

Detection of eluting peptides is performed using both UV at 214 nm (3nl UV cell, Thermo Fisher Scientific, Germering, Germany) and mass spectrometry (Q-Exactive Plus, Thermo Fisher Scientific, Bremen, Germany). The following separation gradients were applied:

| Time (min) | Flow (nl/min) | %A | %B |
| --- | --- | --- | --- |
| 0 | 800 | 95 | 5 |
| 10 | 800 | 95 | 5 |
| 11 | 600 | 97 | 5 |
| 100 | 600 | 75 | 25 |
| 150 | 600 | 40 | 60 |
| 151 | 600 | 10 | 90 |
| 160 | 800 | 10 | 90 |
| 161 | 800 | 95 | 5 |
| 240 | 800 | 95 | 5 |

Mass spectrometry analysis was performed using a Q-Exactive Orbitrap Plus equipped with the Flex nano-ESI source and stainless-steel emitter needle (20 µm ID x 10 µm tip ID). The emitter voltage was set to 3.1 kV, the scan range was 200-2000 m/z. Full MS resolution was set to 70000, AGC target to 3E6, and maximum injection time was set to 50 ms. For the MS/MS analysis, the mass resolution was set to 35.000, AGC target to 1E5, and the maximum injection time to 120 ms. The isolation width for MS/MS was set to m/z 1.5, and top 15 ions were selected for fragmentation, single charged ions and ions bearing a charge higher than +7 were excluded from MS/MS. Dynamic exclusion time was set to 20 seconds. The NCE was set to 30.

Raw MS Data were searched against the human SwissProt protein database (Version November 2020) with Proteome Discoverer 2.4 (ThermoFisher Scientific, Bremen, Germany) using the following parameters: Taxonomy: Homo sapiens; Modifications: carbamidomethyl on C as fixed modification, carboxymethylation on M as a variable modification; Peptide tolerance was set to 10 ppm and the MS/MS tolerance to 0.05Da; Trypsin was selected as the enzyme used and two missed cleavages were allowed; False discovery rate (FDR) was set to 1% and the decoy database search was used for estimating the FDR.

This search yielded >1383 proteins in sEV of melanoma cell lines and >288 proteins in sEV of human plasma samples. Differences in detection frequencies as a proxy for differential exosomal protein composition between groups were calculated and a difference cut-off was set at 50%. Data visualization was performed using the ClustVis tool (18). No centering or scaling was applied to data presented in PCA plots and heatmaps. The mass spectrometry proteomics data have been deposited to the ProteomeXchange Consortium via the PRIDE (19) partner repository with the dataset identifier PXD025475.

## Results & Discussion

In DEL-1 mRNA positive lines only negligible amounts of DEL-1 protein were detectable in lysates and supernatants. In contrast, robust protein expression was seen in sEV derived from wild-type cells and even more from sEV of DEL-1 overexpressing cells (Figure 1a, panel to the left; examples for other DEL-1 positive and negative cell lines are shown in Supp. Fig. 1). A similar secretion pathway for DEL-1 was described in mesothelioma cells (20). The predicted molecular size of DEL-1 is 52kDa; we detected a band at 65kDa, which corresponded to the size of recombinant DEL-1 expressed in Chinese hamster ovary cells (R&D Systems), indicating glycosylation.

**Figure 1.**
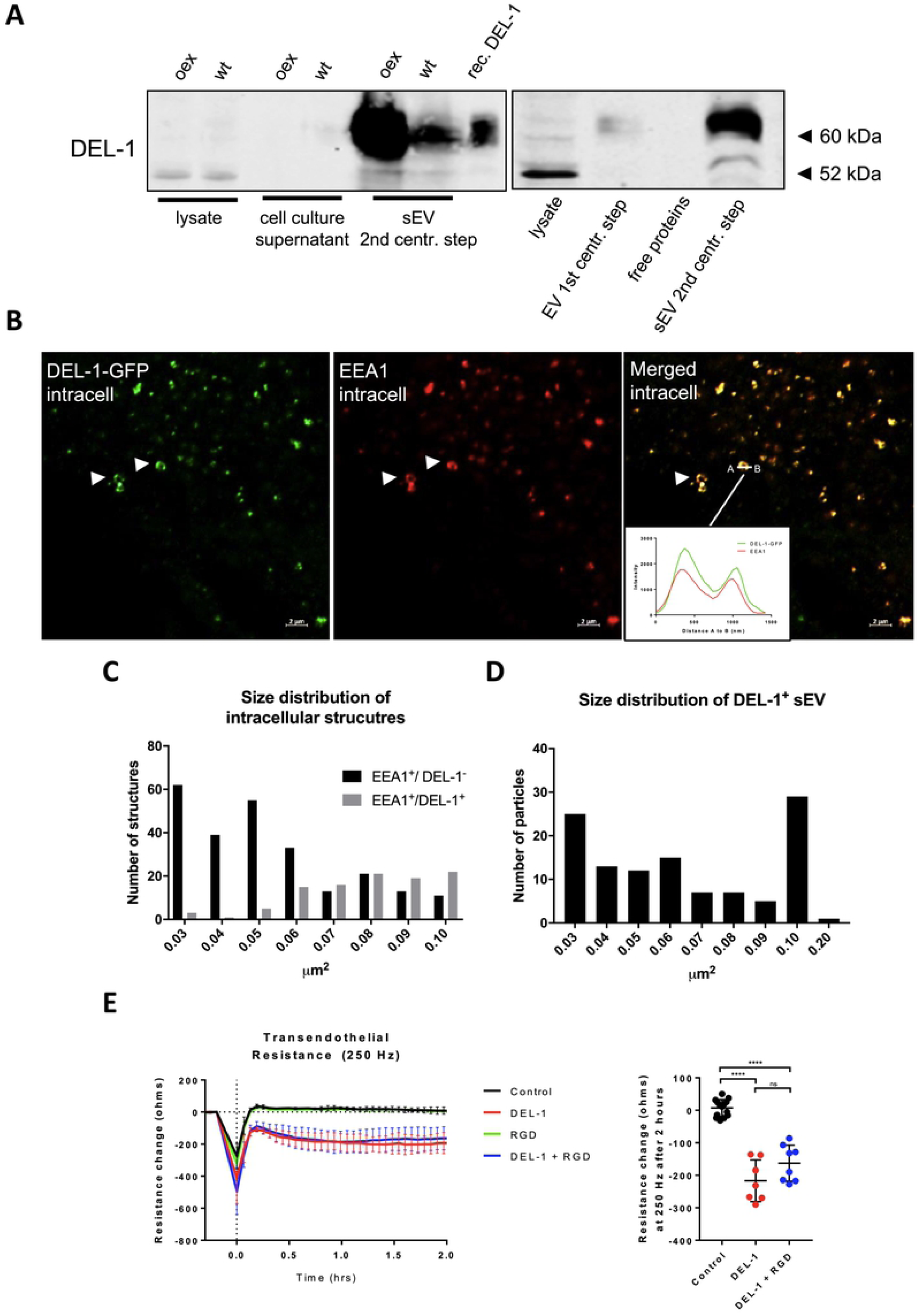
Intracellularly, DEL-1 is localized into endosomal structures, is secreted via sEV and causes endothelial barrier disruption in vitro. a) Left panel: Western Blot analysis for DEL-1 in whole cell lysates, culture supernatants and sEV isolated by a 2-step ultracentrifugation protocol in MCMDLN wildtype and DEL-1 overexpressing cells, recombinant DEL-1 is used as a positive control. Right panel: Western Blot analysis for DEL-1 in microvesicles collected after the 1^st^ ultracentrifugation step for 30 minutes at 18,000xg, and in sEV after the 2^nd^ ultracentrifugation step for 120 minutes at 100,000xg. The nature of the 52kD band in whole cell lysates remains open, but potentially corresponds to the predicted molecular size of 52kD of non-glycosylated DEL-1 (representative example of 3 repeats). “oex” denotes overexpression, “wt” denotes wild type b) Immunofluorescence of a GFP-tagged DEL-1 construct (green) transiently transfected into VM-14 melanoma cells and stained with EEA1 antibody (red). Co-localization is shown in yellow. Arrows indicate early endosomes. Insert shows a histogram of the quantified intensity profile between point A and B to confirm co-localization of DEL-1-GFP and EEA1 (representative example of 2 repeats). c) Size distribution of EEA1 positive/DEL-1 negative, EEA1/DEL-1 double positive intracellular structures as measured as area in µm^2^. d) Size distribution of DEL-1-GFP positive sEV isolated from cell culture supernatant by 2-step differential ultracentrifugation and analyzed by immunofluorescence analysis as measured as area in µm^2^. e) ECIS assay of a confluent endothelial monolayer treated with 10 µg/ml of recombinant DEL-1, 35 µg/ml RGD peptide, a combination thereof or PBS as a control. Transendothelial electrical resistance at 250 Hz was monitored for 2 hours and displayed as resistance change in ohms. Values are shown as mean ± SD. Statistical significance of the resistance change after 2 hours was calculated using one-way ANOVA; ns not significant, **** p <0.0001, n= ≧ 6.

To analyze if DEL-1 is released via an endosomal pathway, we assessed its subcellular distribution prior to its release into the extracellular space. VM14 melanoma cells (8), which do not release DEL-1-positive sEV (Supp. Fig. 1), were transduced with a DEL-1-GFP fusion construct and stained with an anti-early endosome antigen 1 (EEA1) antibody. EEA1 is an effector protein bound to the endosomal membrane (21). Confocal microscopy revealed that in the cytoplasm, DEL-1-GFP forms ring-like structures with a size of up to 1µm, which co-localized with EEA1 (Figure 1b and insert). Size determination of fluorescently labelled structures revealed that the majority of the EEA1 positive but DEL-1 negative intracellular structures were small (below 0.06 µm^2^), whereas EEA1 and DEL-1 double-positive structures were > 0.06 µm^2^ (Figure 1c), possibly corresponding to exosomal cargo loading (22).

EVs are heterogeneous in size. A standard differential ultracentrifugation protocol was used to separate microvesicles (after the 1^st^ ultracentrifugation step for 30 minutes at 18,000xg) from sEV (after the 2^nd^ ultracentrifugation step for 120 minutes at 100,000xg) from conditioned cell culture medium (23). Western blot analysis revealed that DEL-1 is mainly detected in sEV after the 2^nd^ ultracentrifugation step (containing exosomes) with small amounts within microvesicles and no protein is detectable in cell culture supernatants (Figure 1a, panel to the right). Although our characterization of the subpopulation of EVs do not fully meet criteria published in a consensus paper (15), size distribution of the DEL-1-GFP positive sEV fraction as determined by laser scanning microscopy was in the range of 0.03-0.09 µm^2^, which is within the size range of exosomes (Figure 1d). Moreover, protein content-based sEV characterization according to the consensus paper (15) revealed that after the 2-step centrifugation protocol melanoma-derived sEV contain most category 1 proteins required for EV characterization (see methods).

Metastatic homing of circulating melanoma cells to distant organ sites involves their attachment to and transmigration through the endothelium (14). These events require endothelial activation via specific adhesion molecules including integrins. Particularly proteins containing the RGD motif are known to activate endothelial integrins, thereby inducing endothelial retraction (24). We have previously shown that melanoma cells expressing DEL-1 mRNA cause endothelial junction disruption in co-culture (8). The ECIS^®^ system was employed to assess the effects of the integrin-binding protein DEL-1 on endothelial integrity (16) and transendothelial resistance at 250 Hz was measured after treatment of a confluent endothelial monolayer with recombinant DEL-1. DEL-1 induced a pronounced disruption of the endothelial barrier (Figure 1e). DEL-1 contains a RGD finger (1), but co-incubation with a RGD peptide insignificantly reduced DEL-1-induced vascular barrier disruption, potentially due to higher binding affinity of DEL-1 (Figure 1e).

To further characterize differences between DEL-1^+^ or DEL-1^−^ sEV, proteomic profiling of sEV from 8 different melanoma lines published previously (8, 13) was performed. Principal component analysis based on differentially expressed proteins detected by mass spectrometry proteomics data revealed two distinct clusters (Figure 2a), clearly differentiating between sEV from proliferative / DEL-1^-^ and invasive / DEL-1^+^ cells. Detection frequencies of these proteins are shown in Figure 2b. To evaluate, if this can be used to predict patient treatment responses, plasma sEV from 6 melanoma patients undergoing checkpoint inhibitor treatment were collected before and 90 days thereafter. Patient P1 had stable disease, P5 responded and P2, 3, 4, and 6 acquired distant metastasis (Table 1). Unfortunately, from 1 ml of plasma, the sEV yield was poor allowing the identification of 122+35 proteins/patient sample only. DEL-1 was not detectable in any sample.

**Figure 2.**
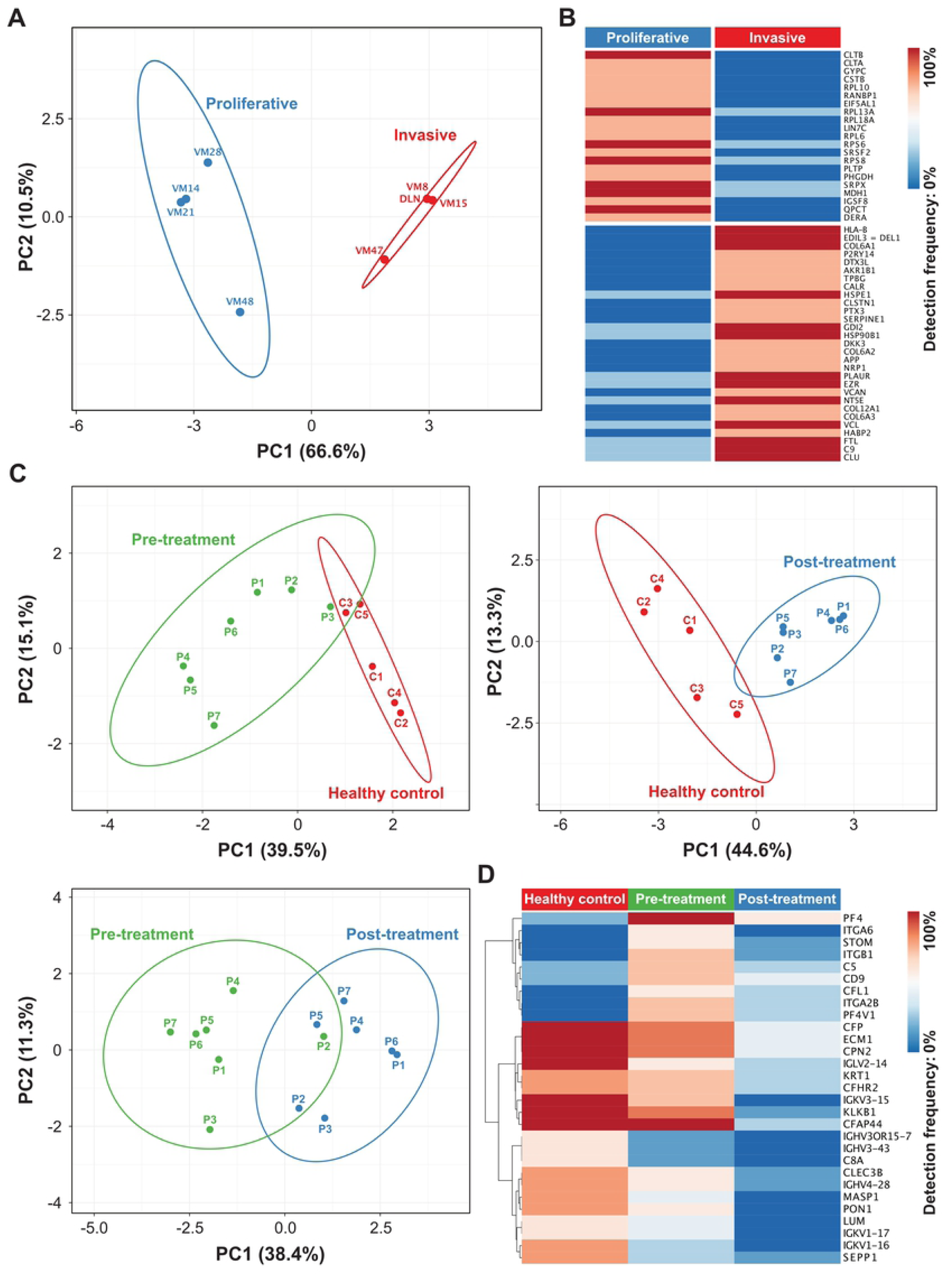
Proteomic profiling of sEV from cultured melanoma cells and plasma sEV from healthy donors and melanoma patients pre- and post-checkpoint inhibitor treatment. a) Principal component analysis (PCA) plot of mass spectrometry data shows qualitative differences in identified proteins between sEV (isolated by a 2-step ultracentrification protocol) from indicated invasive and proliferative cell lines. b) Corresponding heatmap of 50 differentially expressed proteins with 75% detection difference. c) PCA plots show clustering of plasma sEV (isolated by a 2-step ultracentrification protocol) derived from subjects with different health states (healthy, melanoma pre- and post-treatment). d) Corresponding heatmap of 29 differentially expressed proteins between groups with a difference cut-off set at 50%.

The complete data set is published in ProteomeXchange with the identifier PXD025475. **<u>For the reviewer</u>**: Username; PW: UVmO0YGs.

The low protein yield is undoubtedly the limitation of this analysis, and the lack of detectable DEL-1 in patients’ sEV, even in patients with tumor progression. Importantly, the principal component analysis allowed a clear differentiation between controls and patients and to some extent between before and after treatment (Figure 2c, d). Interestingly, the proteomic profile of sEV from patient P5, who showed complete regression, remained in the pre-treatment cluster.

## Conclusions

We provide data that DEL-1, which is thought to be a secreted molecule, is not found in cell culture supernatants but is exported by sEV in melanoma cells. DEL-1^+^ sEV derived from cell culture supernatants co-express many proteins associated with aggressive cancer such as clusterin, vinculin, versican, ezrin, CD87, neuroplin1, SERPINE1, pentraxin3, calreticulin (Figure 2b). The small patient sample size as well as the poor protein yield from plasma-derived sEV is a clear limitation of our study. The apparent difference in protein expression profiles in patients’ sEV pre- and post-treatment as well as compared to healthy controls asks for further studies to analyze its potential to predict treatment responses.

## Data Availability

Data of participating patients and healthy controls and Materials and Methods are described in the Supplementary Materials. Mass spectrometry proteomics data are publicly available via ProteomeXchange with identifier PXD025475. Further data supporting this article are available upon reasonable request.

## Conflict of Interest

The authors declare no conflict of interest.

## Author Contributions

Conceptualization: PP

Patient sample collection: FT

Data Curation: MS, GM, GH

Formal Analysis: PP, MS, GH, MG, JW

Funding Acquisition: none

Investigation: MS, KN, GM, MG

Methodology: MS

Software: GH

Supervision: PP

Validation: PP

Visualization: MS, GH, MG

Writing – Original Draft Preparation: MS, PP

Writing – Reviewing and Editing: MS, PP

## Abbreviations

DEL-1: (developmental endothelial locus-1)
EEA1: (early endosome antigen 1)
sEV: (small extracellular vesicle)
RGD: (Arg-Gly-Asp)

## Notes

### Competing Interest Statement

The authors have declared no competing interest.

